# Modelling perceptual uncertainty in a thermosensory illusion across the lifespan

**DOI:** 10.1101/2025.07.10.664158

**Authors:** Jesper Fischer Ehmsen, Alexandra G. Mitchell, Arthur S. Courtin, Camilla E. Krænge, Cyprien Simonnet, Francesca Fardo

## Abstract

Most studies of perceptual illusions rely on explicit reports alone, offering a limited view into the computations that underlie ambiguous sensory experiences. Here, we introduce a multivariate computational approach to model paradoxical heat sensation (PHS), a thermosensory illusion in which skin cooling evokes sensations of warmth, as an individual-specific process. We tested 75 healthy adults (aged 21–80) using a perceptual decision-making task with stimuli designed to elicit PHS. Binary perceptual choices, response times and confidence ratings were jointly analysed using hierarchical multivariate Bayesian modelling to characterise PHS at both the population and individual levels across the lifespan. We tested two distinct perceptual profiles: “true perceivers”, who experience PHS similarly to a veridical warm percept; and “unsure perceivers”, who perceive PHS as an ambiguous experience. At the group level, behaviour was best explained by the unsure perceiver model, suggesting that PHS is often an uncertain experience, rather than a categorical misperception. At the individual level, however, both profiles were represented, with considerable inter-individual variability in model fit. Older participants were more likely to report PHS and did so at lower levels of thermal contrast, but we observed no correlation between age and perceptual profile. This suggests that while ageing increases the likelihood of experiencing PHS, it does not alter the qualitative nature of its perception. These findings also show how multivariate modelling of perceptual, decisional and metacognitive responses can reveal distinct, subjective profiles and their variation across individuals and age.

## Introduction

Perceptual illusions offer a window into the fundamental principles that shape everyday sensory experiences. Across modalities, they expose how the brain resolves ambiguity, reconciles conflicting cues, and infers meaning from incomplete or noisy inputs. In vision, classic illusions such as the Mach Band and the waterfall effect have highlighted mechanisms like lateral inhibition across the visual system^1–4^. In the somatosensory domain, phenomena such as the thermal grill illusion, the cutaneous rabbit, and referred sensations illustrate how tactile and thermal signals are integrated across time and space to construct coherent percepts^5–11^.

Most empirical work on perceptual illusions has relied on forced-choice tasks, where participants categorise their experience in binary terms (i.e., the illusion is present or absent). While effective in identifying the stimulus conditions that give rise to illusory percepts, such paradigms often overlook the variability and uncertainty of the experience itself. This limitation is particularly evident for illusions that are variably expressed across individuals, occur over relatively long time periods, or are rooted in multisensory integration (e.g., the Thermal Grill or Rubber Hand illusions^12,13^). To address this gap, we introduce a computational model that maps illusory percepts as a multivariate decision process, moving beyond binary categorisation. Our approach jointly models stimulus features with measures of perceptual uncertainty, quantified through trial-wise response times and confidence ratings, to characterise how individuals experience a thermosensory illusion known as Paradoxical Heat Sensation (PHS).

PHS refers to the illusory experience of warmth during cooling, generated by temporally alternating warm and cold temperatures on the skin^14,15^. It is often described as an unusual experience and is more commonly reported in individuals with thermosensory deficits, including those associated with ageing and nerve damage^16–22^. The probability of reporting PHS increases when the contrast between successive warm and cold temperatures is high, in young, healthy individuals^23^. Whilst this finding provides a critical insight into the role of thermal contrast in PHS, it offers only a limited understanding into the perceptual experience itself, including ambiguity, subjective uncertainty and variability across individuals^20,21,24^. For example, a report of warmth during cooling could reflect a true perceptual shift, or instead reflect a lack of clarity about the thermal quality with “warm” being the closest descriptor. Without additional behavioural markers to characterise this uncertainty, such distinctions remain inaccessible.

To quantify illusory perception, we analysed perceptual binary choices (“cold” or “warm”) alongside the associated response times and confidence ratings. These trial-wise measures serve as behavioural proxies for decision latency and perceptual certainty. Incorporating them into a multivariate computational model allowed us to infer how thermosensory input is interpreted under uncertainty and how different perceptual profiles manifest across individuals.

We generated and tested two theoretical perceptual profiles to describe the PHS experience, and its link to thermal contrast: the “true perceiver” and the “unsure perceiver”. The “true perceiver” interprets PHS as a true illusion, where the perceiver experiences PHS as a sensation akin to veridical warmth. In this profile, perceptual uncertainty peaks near the PHS threshold, where the probability of experiencing PHS is around 50%. This is reflected by slower response times and lower confidence at thermal contrasts near the PHS threshold. In contrast, the “unsure perceiver” experiences PHS as an ambiguous sensation that is neither distinctly warm nor cold. For this profile, uncertainty increases monotonically with thermal contrast, peaking at high contrast levels where the probability of reporting PHS is highest.

To evaluate these accounts, we implemented two computational models, each formalising one of the perceptual profiles, and tested their ability to quantify the perceptual uncertainty associated with PHS in our entire sample. The winning model was used to jointly assess the effects of thermal contrast and ageing on PHS, response time and confidence. Additionally, we examined the extent to which each model best explains the response profile of each individual. Crucially, this includes not only those who report PHS but also “non-responders” who never report the illusion. Our approach provides an in-depth characterisation of PHS experience, distinguishing between perceptual content, response dynamics and subjective uncertainty, in relation to stimulus properties and ageing, that is broadly applicable beyond thermosensation. The distinction between true and unsure perceivers may generalise to other domains involving ambiguous or unstable percepts, such as visual motion illusions, multisensory integration phenomena or clinical distortions of body perception. As such, this model provides a powerful tool for characterising individual differences in the construction of conscious sensory experience.

## Results

### Ageing, thermal contrast and PHS

To characterise the experience of thermosensory illusions across the lifespan, we tested a perceptual decision-making paradigm in 75 participants (ages 21-80). Participants experienced dynamic thermal stimulation consisting of alternating warming and cooling temperature ramps. Participants were asked to indicate when they noticed a change in temperature, during the cooling phase, and whether it felt cold (i.e., veridical sensation) or warm (i.e., PHS) (Fig. 1). The magnitude of the difference between alternating warm and cold temperatures, referred to as thermal contrast, was manipulated across trials by varying the peak temperature of the warming phase (Fig. 1 A & B), spanning from innocuous to mildly noxious levels. Although all perceptual decisions were made during cooling, participants were unaware of the underlying direction of temperature change. This design dissociated physical input from perceptual interpretation, allowing us to probe how subjective thermal experiences, veridical or illusory, emerge from alternating thermal stimulation.

**Figure 1:**
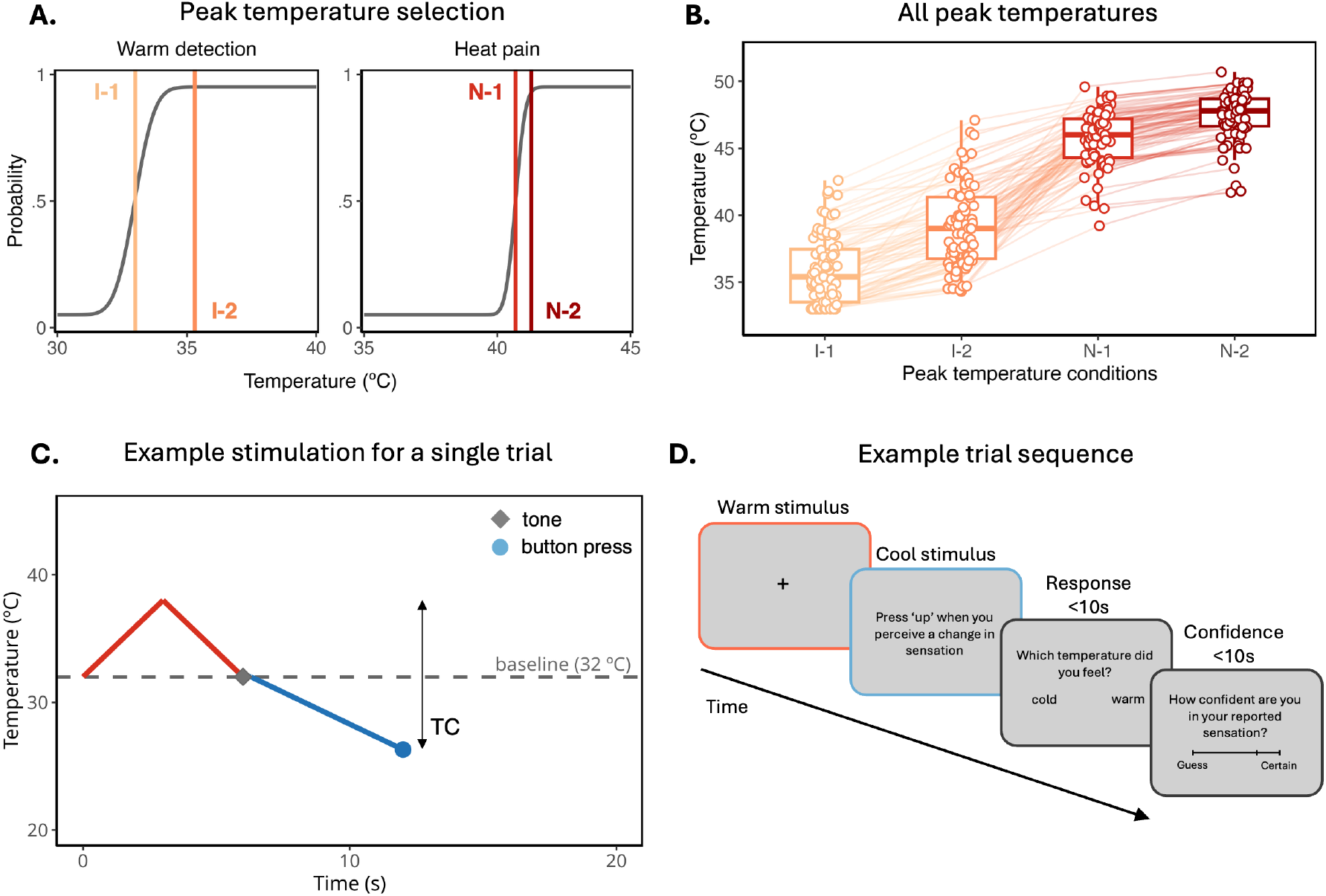
Task overview and thermal stimulation parameters. (A) Example psychometric functions used to select participant-specific peak temperature, derived from prior thresholding. Vertical lines represent the four peak temperatures used to manipulate thermal contrast across trials. Two peak temperatures in the innocuous range (I) were derived from warm detection thresholds and slopes, while two peak temperatures in the noxious range (N) were derived from heat pain thresholds and slopes. (B) Selected peak temperatures for all participants. (C) Time course of stimulation for a single representative trial, showing the alternating warm (red) and cold (blue) temperature ramps. The magnitude of thermal contrast (TC) was defined as the difference between peak warm and cool temperatures, divided by the total thermal range (Eq. 1). (D) Trial sequence. Following a brief warming phase, a cooling stimulus was delivered. Participants were instructed to press a key when they detected a change in temperature, then report whether the change felt cold (veridical) or warm (PHS), followed by a confidence rating. For illustrative purposes, the colour of the screen border represents whether the probe was above (red), below (blue) or at baseline temperature (grey).

A total of 64/75 participants (85%) reported experiencing a PHS at least once. Out of all experimental trials (56 per participant), 22% were experienced as PHS. Both the rate of reported PHS and PHS prevalence increased with increasing peak temperature (Fig. S1). Figure 2 provides a comprehensive overview of the relationship between thermal contrast, peak temperature and age on PHS, response time and confidence.

**Figure 2:**
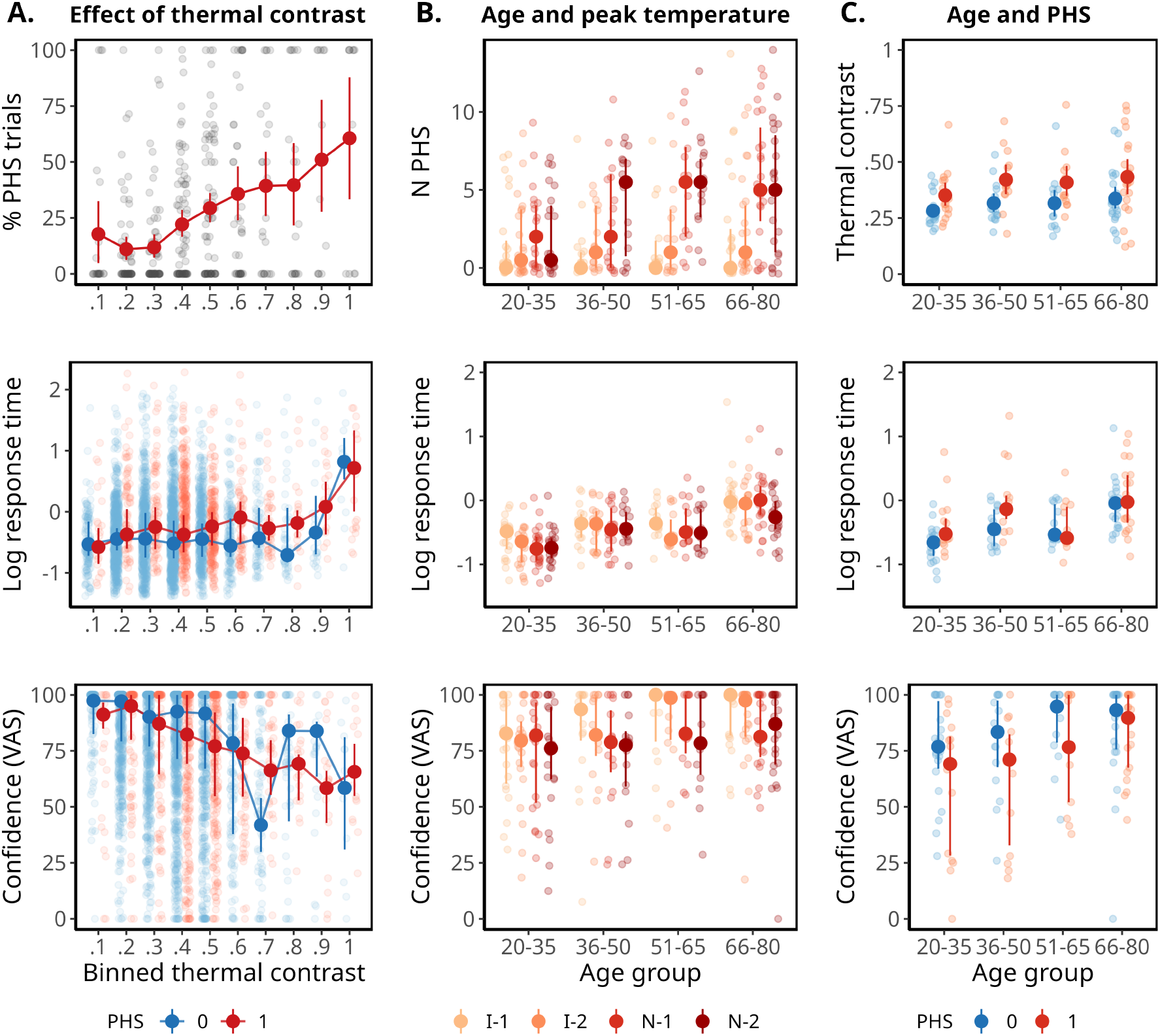
Effects of thermal contrast, peak temperature and age on PHS, response time and confidence. (A) Percentage of PHS trials, log-transformed response time and confidence VAS ratings plotted against binned thermal contrast levels. (B) Relationship between age group and responses grouped by peak temperature condition (I-1, I-2, N-1 and N-2). (C) Effects of age and PHS on thermal contrast, response time and confidence ratings. Small dots show individual participant means (% PHS and thermal contrast) or medians (N PHS; response time and confidence); large dots represent group means with 95% confidence intervals (% PHS and thermal contrast) or medians with interquartile ranges (N PHS, response time and confidence). Binning and grouping are used for visualisation only; all variables were modelled as continuous in the analyses.

### A computational account of PHS

To test our hypotheses about the experience of PHS within our sample, we developed two multivariate models that jointly characterise the relationship between thermal contrast, PHS, response time, and confidence. Each model is based around two possible PHS profiles - true perceiver and unsure perceiver - that incorporate binary perceptual classifications (“cold” or “warm”), choice response time and confidence rating (Fig. 3). Both models quantify the relationship between PHS and thermal contrast as a logistic psychometric function, where the probability of PHS is expected to increase monotonically with increasing thermal contrast (Eq. 2). Confidence ratings and response times are then modelled as functions of PHS probability, allowing us to test different patterns of association between these indices of perceptual uncertainty and the likelihood of experiencing PHS.

**Figure 3:**
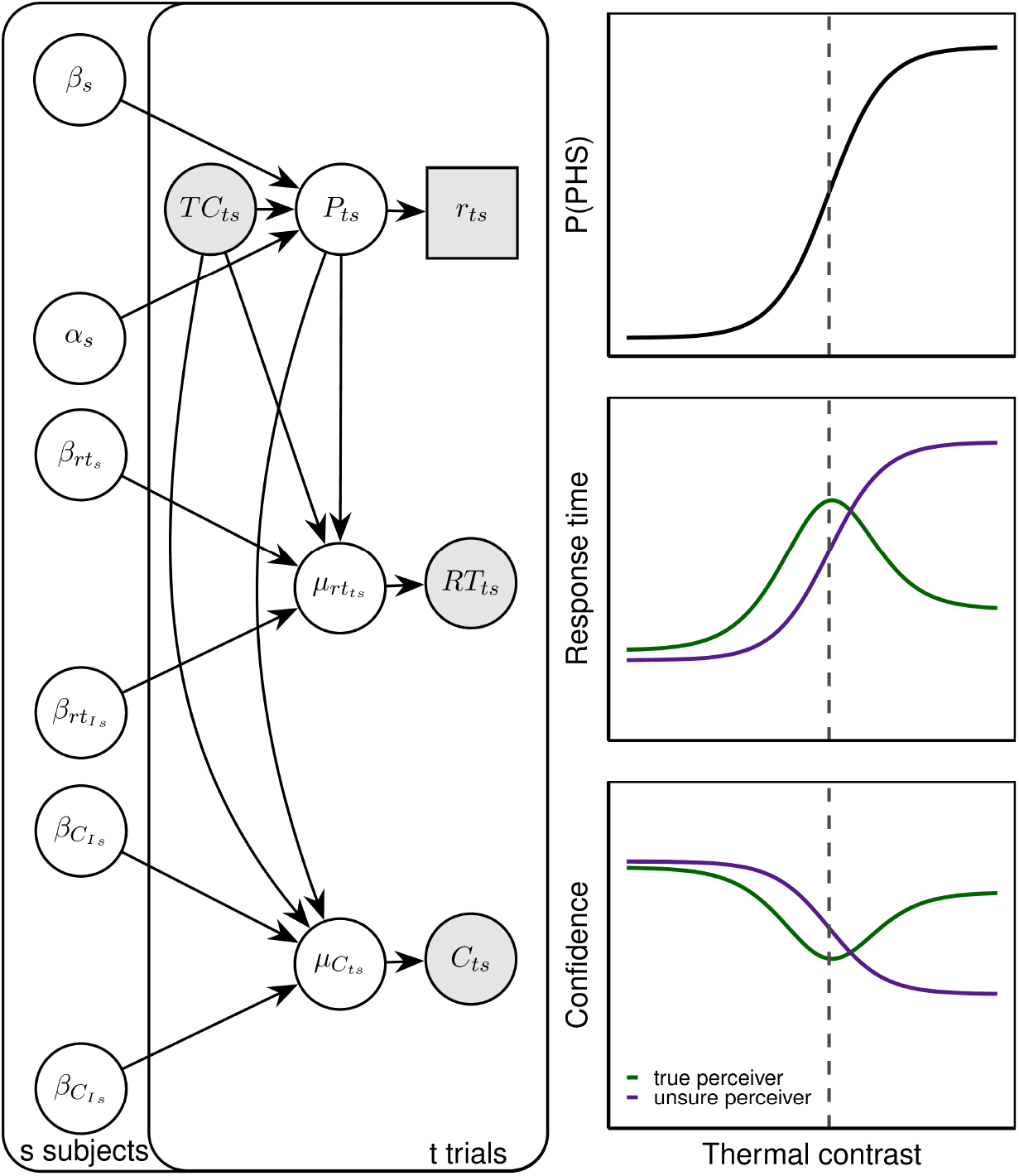
A multivariate model of PHS, response time and confidence. Left: Plate notation illustrating the generative structure of the hierarchical model for a single subject. The model jointly estimates perceptual outcomes (PHS reports, *P*_*ts*_), response times (*RT*_*ts*_), and confidence ratings (*C*_*ts*_) as a function of thermal contrast (*T C*_*ts*_) and subject-specific parameters. For clarity, nuisance parameters (e.g., residual variance) and hierarchical population structure are omitted from the diagram. Right: Simulated model outputs showing predicted effects of thermal contrast on PHS probability, response time and confidence under two model variants: true perceiver (green) and unsure perceiver (purple). The true perceiver model assumes that response time and confidence are linked to decision uncertainty (entropy), with maximum uncertainty at the perceptual boundary. The unsure perceiver model assumes that response time and confidence scale directly with the probability of experiencing PHS. These models provide alternative mechanistic accounts of how individuals resolve perceptual ambiguity in response to alternating thermosensory input.

The true perceiver model (model 1) assumes that perceptual uncertainty in PHS arises from the difficulty of making a categorical choice between “cold” and “warm”, rather than from the experience of PHS itself. This model predicts the highest uncertainty where the stimulus is at the perceptual boundary between cold and warm, and the highest certainty at the points at which the sensation is either clearly cold, or clearly warm. Uncertainty is reflected by the slowest response times and lowest confidence which occurs at thermal contrast values associated with 50% probability of experiencing a PHS. This pattern aligns with the concept of a “true” PHS responder as an individual who experiences a clear and definite warm sensation in response to a cold stimulus.

The unsure perceiver model (model 2) assumes that perceptual uncertainty does not arise from the difficulty of making a categorical choice between “cold” and “warm”, but from the aberrant and unusual experience of PHS. Therefore, uncertainty increases with the probability of experiencing PHS itself, with slowest response times and lowest confidence ratings expected at the highest thermal contrast values. This profile is called the unsure perceiver as it reflects increased uncertainty at thermal contrast values where participants more frequently report PHS, reflected in slower response times and lower confidence ratings.

Our two multivariate models were conducted in Stan^25^ using [pre-registered](https://osf.io/t5nmd) priors and sampling parameters (see Supplementary Materials). We found that the unsure perceiver model provided a significantly better out of sample fit than the true perceiver model with an expected log predictive density difference of -29.67 ± 10.19, thus providing an improvement in model fit by -2.91 standard errors of the difference.

### Both thermal contrast and the probability of PHS increase in older adults

We used our winning model, the unsure perceiver model (model 2), to test how ageing and thermal contrast influences the likelihood of PHS and the associated behavioural markers: response time and confidence. Figure 4 presents data from four example participants. Data from all participants is presented in the Supplementary Materials (Figs. S2-S4).

**Figure 4:**
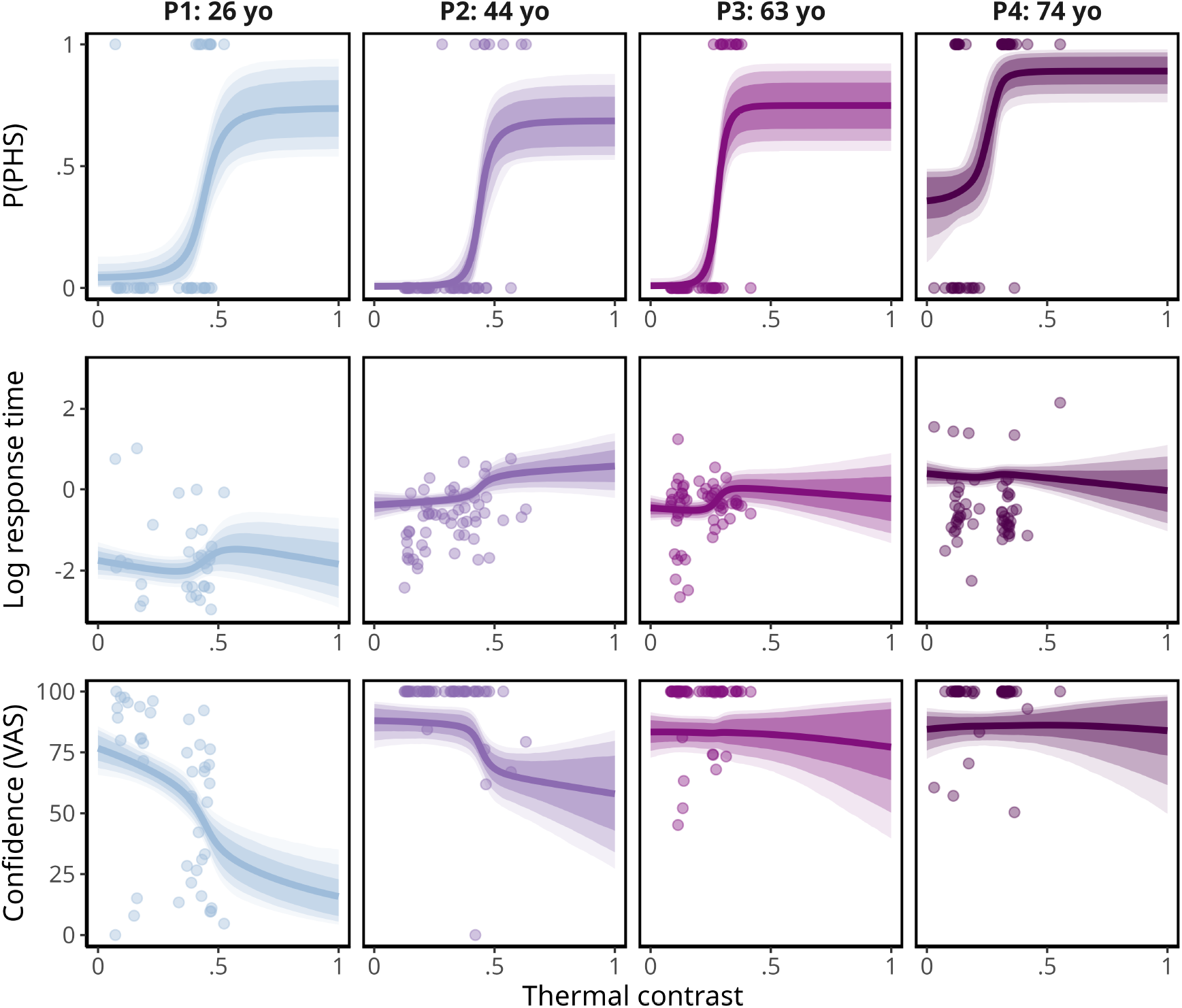
The modelled change in PHS probability, response time and confidence rating as a function of thermal contrast for four example participants (P1 - P4). Each column shows data from a single participant, ordered from youngest to oldest. Rows represent posterior estimates for the probability of reporting PHS, log-transformed response time, and confidence ratings as a function of thermal contrast. Points indicate trial-wise data; thick lines show posterior means from our multivariate model; shaded areas represent 90%, 80% and 60% credible intervals (from light to dark). These examples illustrate individual variability in PHS perception and uncertainty across age.

At the group-level, we found that the probability of experiencing PHS is higher in older adults (Fig. 5). For each additional year of age, the thermal contrast required to reach a 50% probability of PHS decreased by -0.68 (Δ*µ*_*phs*_ = -0.68, 95% highest density interval [HDI] [-1.39, 0], Posterior probability (*P*) = .02), indicating increased PHS sensitivity with age. As expected, older adults also showed slower response times (Δ*µ*_*rt*_ = 0.02, 95% HDI [0.01, 0.02], *P* = 1). Additionally, increased probability of PHS was associated with slower response times (*β*_1*rt*_ = 1.48, [0.95, 2.09], *P* = 1). Increasing age was associated with higher mean confidence ratings (Δ*µ*_*conf*_ = 0.01, 95% HDI [0, 0.02], *P* = .97). However, confidence ratings decreased as PHS probability increased (*β*_1*conf*_ = -1.25, 95% HDI [-1.87, -0.66], *P* = 0).

**Figure 5:**
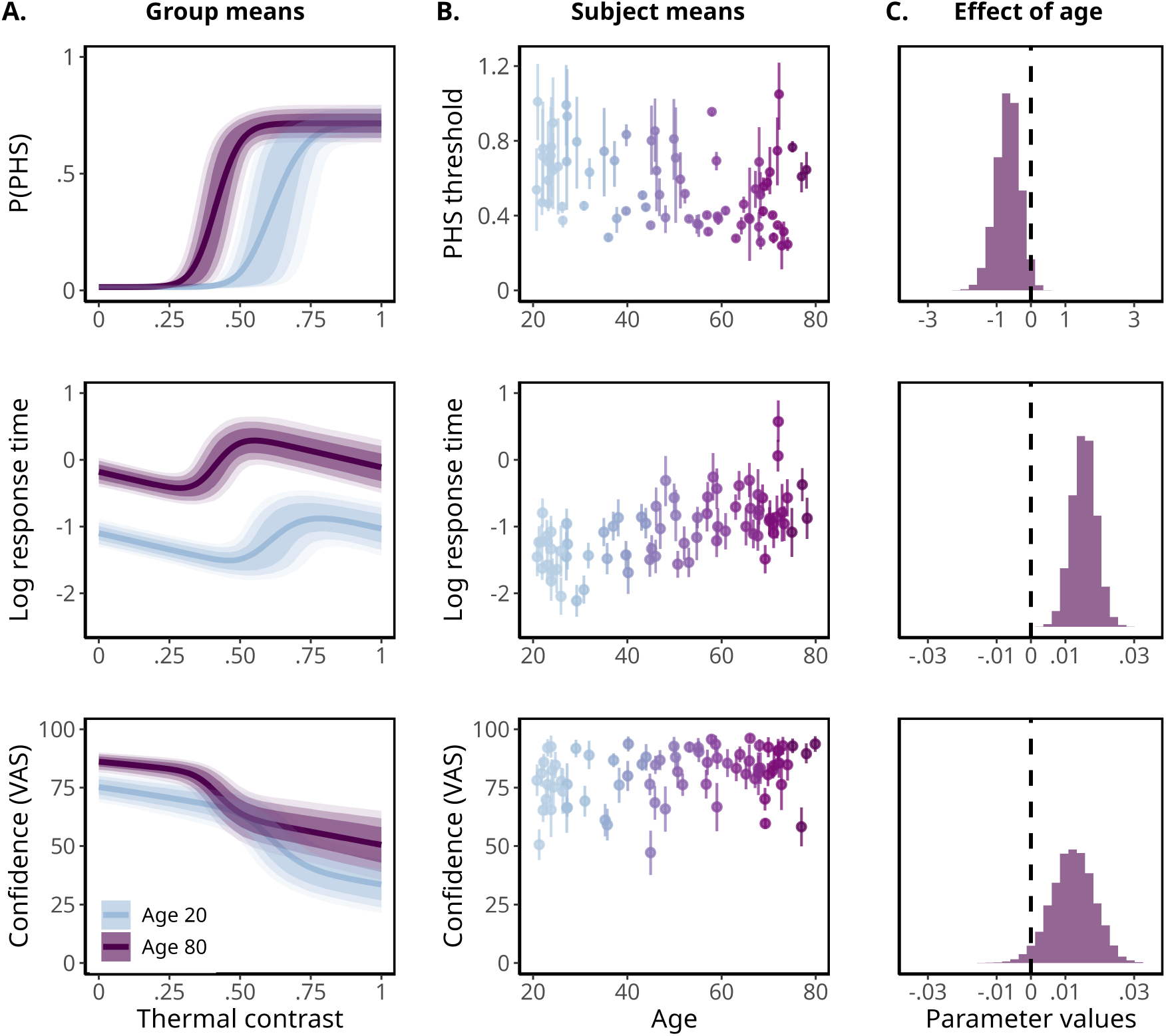
Group-level estimates for the winning model (unsure perceiver) and age-related changes in perceptual response. (A) Posterior predictions from the best-fitting model (unsure perceiver) showing group-level estimates for the effect of thermal contrast on probability of PHS, response time and confidence ratings. Curves are plotted for ages 20 and 80, corresponding to the lower and upper bounds of the sample age range. Solid lines indicate posterior means, shaded regions show 90, 80 and 60% credible intervals. (B) Individual-level estimates of PHS threshold, mean response time and mean confidence ratings plotted by age. Points represent posterior means per participant and error bars indicate the 68% credible interval. (C) Posterior distributions of the estimated effect of age on each parameter, quantifying the average change per year (displayed as a percentage change for PHS threshold). Distributions reflect both the magnitude and direction of age-related shifts for PHS probability, response time and confidence.

To better quantify the magnitude of the effects of thermal contrast and probability of PHS on response time and confidence ratings, we computed conditional mean differences. When the probability of PHS was fixed to be 0, the mean difference in response time between minimum (0) and maximum (1) thermal contrast was -0.32 s (95% HDI [-0.44, -0.2]), indicating that higher contrast leads to faster responses. The corresponding mean change in confidence was -18 points (95% HDI [-35, -3]), indicating a substantial decrease in confidence with thermal contrast. Conversely, when thermal contrast was held constant at 0, increasing the probability of PHS from its minimum to maximum was associated with a mean increase in response time of 1.01 s (95% HDI [0.4, 1.73]), indicating slower responses as PHS probability increased. Confidence ratings simultaneously decreased by -17 points (95% HDI [-28, -8]), highlighting lower confidence as PHS probability increased.

### Individual differences in the experience of PHS

Our modelling approach allowed us to examine individual variability in the experience of PHS. We expected that some participants would conform to the “true perceiver” profile, characterised by consistent and confident PHS reports, whereas others would be aligned with the “unsure perceiver” profile. To quantify individual model preference, we fit non-hierarchical models to each participant and compared ELPD values for the two profiles at the individual level. We defined the ELPD difference such that negative values indicated a preference for the true perceiver model, and positive values for the unsure perceiver model. Although the unsure perceiver model provided the best fit at the group level, model comparison at the individual level revealed substantial heterogeneity in PHS response patterns (Fig. 6A). Some participants demonstrated responses consistent with the true perceiver, where uncertainty was maximal around the PHS threshold, whereas others showed increasing uncertainty as PHS probability increased, reflecting distinct perceptual profiles within the sample. We further examined whether model preference varied with age, and found no significant correlation between age and model preference (*ρ* = .13, 95% CI [-.09, .33], *p* =.87) (Fig. 6B).

**Figure 6:**
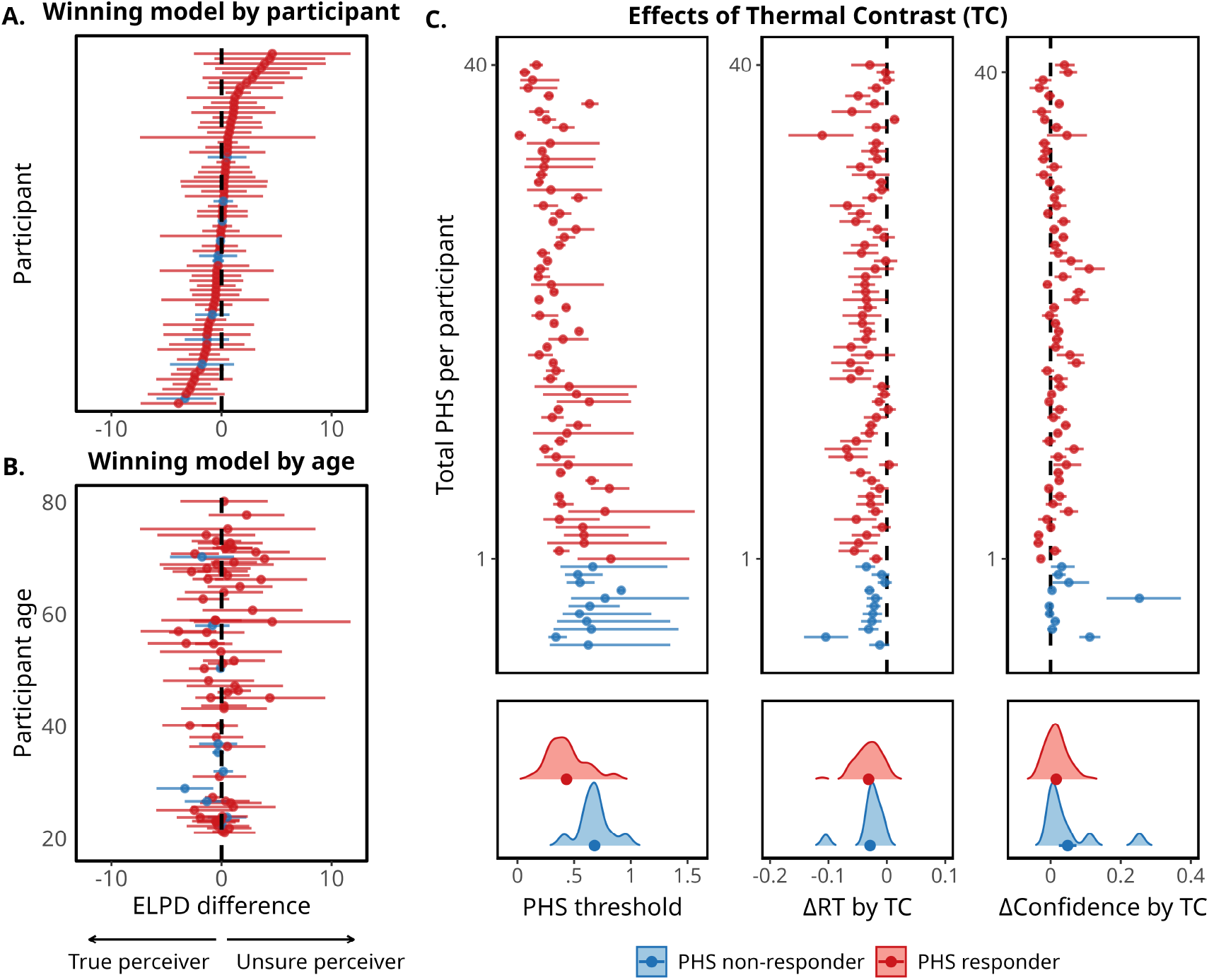
Individual perceptual profiles of PHS. (A) ELPD difference values from approximate leave-one-out cross validation non-hierarchical model comparison for each participant. Negative values indicate a better fit for model 1 (true perceiver profile), whilst positive values favour model 2 (unsure perceiver profile). Values are sorted by model fit (A) or by participant age (B). (C) Parameter estimates for PHS threshold, the slope of response time and confidence by thermal contrast, ordered by each participant’s total number of PHS reports. Non-responders are indicated in blue. Across all panels, error bars show 95% confidence intervals, and density plots show the distribution, mean and standard error of mean estimates for PHS responders (red) and non-responders (blue). For visualisation purposes, an outlier was removed from the far confidence rating display in (C) where estimated confidence slope was 1.1, the inclusion of which compressed the scale limiting visibility for the rest of the sample. Fig. S5 shows the same figure with the inclusion of this participant.

### Non-responders

A subset of participants (11 out of 75) were identified as PHS non-responders. By definition, these participants reported no PHS, resulting in an almost flat psychometric function with near zero probability of PHS. Although they consistently reported all temperature changes as cold, it is possible that their experience included ambiguous sensations, neither cold nor warm, constrained by the forced-choice paradigm. To assess whether the highest contrast trials elicited a latent, unreported PHS experience, we examined whether thermal contrast influenced their response times and confidence ratings. If a subthreshold increase in PHS was present, it could manifest as systematic changes in these behavioural measures, despite the probability of PHS being fixed at zero. Specifically, this would be reflected in either a stronger-than-expected effect of thermal contrast on response times relative to responders or a corresponding decrease in confidence. At the group-level, we observed faster response times and lower confidence with increasing thermal contrast (Fig. 5A). At the subject-level, posterior estimates confirmed this same pattern in response times for both responders and non-responders (Fig. 6C), while confidence remained largely unaffected. This indicates that, although response times were sensitive to thermal contrast, there was no evidence that higher contrast reduced uncertainty for non-responders.

## Discussion

Paradoxical heat sensation (PHS) occurs when alternating warm and cool stimuli lead to a percept that is reported as warm^14,26^, yet previous research offers little insight into how individuals experience PHS. To achieve this, we developed and tested a computational model that jointly estimates the probability of reporting PHS and the associated perceptual uncertainty. Our model incorporates binary perceptual reports with response times and confidence ratings, allowing us to quantify PHS as a perceptual state shaped by thermal contrast, age-related changes, and individual differences in thermosensory processing. This extends beyond current psychophysical models that are able to estimate perceptual probability but do not account for uncertainty associated with the reported sensation.

We formalised two perceptual profiles that differ in how uncertainty varies with the probability of PHS. In the true perceiver profile, uncertainty is lowest when the probability of PHS is near zero or one and peaks at the 50% threshold. This models PHS as an experience akin to true warmth perception. In the unsure perceiver profile, uncertainty increases with the probability of PHS and reaches its highest point when PHS is most likely. This models PHS as a perceptual state that remains ambiguous or unusual and might not clearly correspond to a perceptual category. At the group level, model comparisons showed the unsure perceiver profile provided a better fit, indicating that for most participants, PHS is associated with increased decisional uncertainty. This finding challenges the view that PHS is typically a clear but incorrect perception of warmth^14,26^ as, for many in our sample, PHS appears to be a perceptual state that may resemble warm perception but is accompanied by uncertainty about how to interpret it. This aligns with the idea that some illusory experiences reflect ambiguity in sensory processing rather than a categorical misperception. For example, similar patterns have been observed in other thermal illusions, such as the thermal grill, where sensations do not always map neatly onto established sensory labels^6,12,27^.

Group-level results based on the unsure perceiver model indicated that PHS probability increased with increasing thermal contrast, in line with previous results^23^. In addition, older adults reported PHS more frequently and at lower levels of thermal contrast. Peak temperatures were individually calibrated to account for differences in thermal sensitivity, therefore this age-related increase in PHS cannot be explained by general sensory decline alone. Instead, it suggests that ageing affects how ambiguous thermal input is processed, independent of perceptual thresholds. This can be interpreted in light of previous work in individuals with spinal cord injury affecting thermosensory pathways^28^. In regions where only one thermal modality remained intact, the quality of non-painful thermal sensation was determined not by the stimulus itself, but by the preserved modality. For example, cold stimuli were perceived as warm in regions where only warm sensation was intact, and vice versa. These paradoxical percepts reflect cross-modality misattribution, in which noisy or absent sensory inputs from one modality are interpreted through the preserved modality. We propose that a similar process may be at play in healthy ageing. Ageing disproportionately affects cold sensitivity, with elevated detection thresholds and shallower psychometric slopes compared to warm detection^29^. Our results that older adults exhibited increased PHS rates and experienced PHS at lower thermal contrast levels, are consistent with the idea that weakened cold input leads to percept biased toward the warm modality^30^. In this view, PHS in ageing may represent a perceptual inference biased by the reliability of the most preserved thermal modality.

In addition to the group-level findings, we tested individual differences in PHS response. Although the majority were better described by the unsure perceiver profile, a subset of participants were better fit by the true perceiver profile, with similar distributions of the two profiles across age. These results indicate the presence of two distinct perceptual profiles highlighting that PHS does not represent a uniform perceptual experience across individuals. This pattern is consistent with findings from other sensory domains, where illusions often show substantial inter-individual variability, including in vision^31,32^ and multisensory illusions such as the rubber hand and other bodily illusions^13,33,34^. Crucially, the distribution of these profiles was consistent across the lifespan, with no correlation between individual model fit and age. This indicates that the age-related increase in PHS is not explained by one perceptual profile becoming more common in older adults. The mechanism underlying the age effect reflects a general increase in the likelihood of the sensation occurring, rather than a change in how it is experienced.

Finally, we identified 86% of participants to be PHS responders, with a subset of participants (11 out of 75) not reporting PHS under any condition. These individuals showed no indication of latent PHS responses, even at the highest levels of thermal contrast. This suggests that these participants experienced the stimuli as veridical cold, without generating the ambiguous percept characteristic of PHS. The absence of any behavioural markers indicative of subthreshold PHS, such as changes in response time or confidence, further supports this interpretation. These results indicate that approximately 15% of individuals may not experience paradoxical heat under the conditions tested.

Together, these results show that PHS is not a uniform perceptual phenomenon but varies both in whether it occurs and in how it is experienced across individuals. This variability, including the presence of non-responders, highlights the importance of modelling not only perceptual outcomes but also the uncertainty surrounding them. Our approach provides a principled way to assess how changes in stimulus properties influence both the probability of a percept and its associated uncertainty. Although developed in the context of thermosensation, the method is applicable to a wide range of sensory and cognitive processes. It may be particularly relevant for more variable and bistable illusions, where modelling behavioural markers of uncertainty can offer new insights into both the nature of these experiences and sources of individual variability. More broadly, this framework has potential applications in clinical contexts, including conditions such as chronic or spontaneous pain, where altered perceptual uncertainty may contribute to the emergence or maintenance of symptoms.

### Strengths and Limitations

This study combined a computational model with a controlled experimental paradigm designed to quantify both the probability of PHS and the associated perceptual uncertainty. A key strength was the use of individually calibrated peak temperatures, which accounted for differences in thermal sensitivity across participants and age. This design ensured that age-related effects could be attributed to changes in perceptual processing rather than baseline differences in thermosensory detection thresholds. The large number of trials per participant also improved the reliability of parameter estimates and likely contributed to the high prevalence of PHS observed in this sample (∼85%), which exceeds rates reported in previous studies^24,35^.

Within our models, we have assumed conditional independence of the response variables (binary choices, response time and confidence ratings) and that any shared dependence is linked to thermal contrast and PHS probability. This simplifying assumption may overlook potential dependencies between measures that are not fully accounted for by the predictors, for example between the response variables themselves. Future work could address this limitation by adopting multivariate frameworks that explicitly model interdependencies among response variables, such as copula-based approaches^36,37^. Finally, whilst we are able to speculate on the mechanisms that might link ageing with increased PHS, our study does not include neurophysiological data (e.g., the degradation of cold input over the lifespan) that can directly support this interpretation. However, we have recently shown that cold perception is particularly susceptible to healthy ageing in the same sample of participants^29^, providing support for our proposed mechanisms. Future research should focus on directly linking the thermosensory effects of ageing with changes to neurophysiology.

### Conclusion

In this study, we applied a multivariate computational model to quantify the experience of an elusive thermosensory illusion, paradoxical heat sensation. The results show that PHS often reflects an ambiguous perceptual state rather than a clear misperception of warmth, with most participants exhibiting high uncertainty when PHS is most likely to occur. Older adults reported PHS more frequently and at lower thermal contrast levels, but this increase was not accompanied by changes in how the experience was qualitatively structured. The presence of two distinct perceptual profiles, consistent across age, demonstrates that PHS is not a uniform perceptual phenomenon but varies meaningfully across individuals in terms of certainty and interpretation. These findings advance the understanding of how ambiguous sensory input is processed and how uncertainty contributes to the experience of perceptual illusions. Critically, the model introduced here offers a general approach for investigating how uncertainty shapes perception, with applications extending beyond thermosensation to other forms of perceptual ambiguity, illusions, and clinical contexts involving altered perceptual experiences.

## Methods

### Study Design

The study employed a single-session, cross-sectional design where participants completed three tasks aimed at assessing thermal sensitivity (detection and pain thresholding), paradoxical heat sensation (PHS), and spatial aspects of thermal perception. In addition, all participants completed survey measures related to handedness, mental health, and pain sensitivity. The experiment took place at the Aarhus University Hospital, Aarhus, Denmark, and was approved by the Region Midtjylland Ethics Committee and adhered to the current version of the Declaration of Helsinki (2013). Participants provided informed consent prior to their participation in the study and were compensated for their participation. This paper focuses on a specific subset of these data, related to the effect of age on PHS which was pre-registered before data analyses.

### Participants and Recruitment Criteria

Inclusion criteria required all participants to be between 20 and 80 years old, with no evidence of neuropathy or pain conditions, and fluent in Danish and/or English. Exclusion criteria included a diagnosed neuropathic or clinical pain condition, a known neurological or psychiatric disorder, pregnancy or lactation, recent jet lag or sleep deprivation, skin diseases such as eczema or psoriasis, a history of alcohol or drug abuse, cannabis use within the previous four weeks, and alcohol consumption within the previous 48 hours. Participants were also excluded for having a condition that may interfere with the study goals (e.g. cognitive decline or a physical disability) or that required them to take medication that is known to affect physiological pathways for either temperature or pain perception. To ensure balanced representation across the adult lifespan, participants were allocated into four age groups: 20-35; 36-50; 51-65; 66-80, where we aimed to recruit between 15 and 25 participants per group.

A total of 84 healthy individuals took part in the study. During data screening, nine participants were excluded due to medication use (e.g., antihypertensive medications). Although these individuals self-identified as physically healthy, such medications may influence physiological responses relevant to temperature perception. Therefore, data from 75 participants (45 F, mean age = 49.71, range = 21–80) were included in the final analyses (see Supplementary Materials for clarification of final numbers).

### Apparatus

All stimuli were delivered using a T11 Thermal Contact Stimulator (QST.Lab, Strasbourg, France), with a total stimulation surface of 90 *mm*^2^ and five independently controlled stimulation zones (one zone 7.4 × 24.2 mm) and were applied to the non-dominant forearm. For each trial two neighbouring stimulation zones, chosen randomly, were either heated or cooled while the remaining three zones were maintained at a predetermined baseline temperature. All experimental tasks were coded in MATLAB R2021a, and presented using Psychtoolbox 3^38,39^.

### Questionnaires

The short-form of the Edinburgh Handedness Inventory (EHI,^40,41^ was used to infer participant handedness. Participants then completed a series of questionnaires to assess pain sensitivity, pain experience in their daily life and mental health factors. Specifically, the State-Trait Inventory for Cognitive and Somatic Anxiety (STICSA), both the current and the general versions^42^, the Pain Sensitivity Questionnaire (PSQ,^43^), the Michigan Neuropathy Screening Instrument (MNSI,^44^) to screen for probable peripheral neuropathy, as well we to gather information on their pain experience and pain sensitivity. Finally, the Patient Health Questionnaire (PHQ-9,^45^) was used as a brief measure of mental health status.

### Detection and Pain Thresholding

Thermal sensitivity was assessed with four brief computerised tasks designed to measure the participants’ ability to detect cooling and warming, as well as their sensitivity to painful cold and heat. These tasks were always performed after the end of the surveys and in the same order: cold detection followed by warm detection on the non-dominant volar forearm, then cold pain followed by heat pain on the dominant volar forearm. For the purposes of this study, only warm detection and heat pain thresholding were used to inform stimulus parameters, therefore we only report on those specific procedures.

A single threshold trial consisted of 1-3 s of stimulation (jittered) with the thermal probe at baseline temperature (30 ºC). After which, the two active stimulation zones increased in temperature at a rate of 2.5°C/s and were maintained at target temperature for 1.5 s. After the 1.5 s plateau, participants were asked “Was the stimulus warm?” for warm detection, or “Was the stimulus burning?” for heat pain, and used the left and right arrow keys to respond either “yes” or “no”. After the response, the active zones decreased back to baseline temperature at a rate of 80 ºC/s during which the participant’s chosen response was displayed for 0.5 s. For all tasks, participants had a maximum of 10 s to respond to each question, using the left and right arrow keys (left - “yes”, right - “no”). The warm detection task consisted of 30 trials and lasted 5 to 7 minutes. The heat pain thresholding task consisted of 50 trials, and was completed within 10 to 15 minutes.

Detection and pain perception psychometric functions (PF) were assessed using the psi method, implemented in the Palamedes toolbox^46^. Psi is a Bayesian adaptive algorithm that iteratively updates a posterior probability distribution of the threshold and slope of a PF, allowing optimal stimulus placement for threshold and slope estimations^47^. This approach ensures that stimulus intensities were calibrated to individual sensitivity, limiting the number of trials at uninformative (never or always detected) or excessively painful intensities. As a result, it reduces the number of trials needed to attain a given level of estimation precision and the risk of habituation or sensitisation. Full details of the psi procedure and adaptive thresholding stimulus selection are available [here](https://doi.org/10.17605/OSF.IO/DVZ94).

Out of the 50 trials used to assess pain sensitivity, the first 10 were familiarisation trials, where the intensity was set using an up-down staircase algorithm rather than psi. This ensured that the participant had some time to stabilise their strategy regarding what they reported as burning. These familiarisation trials were not included in the analyses. The extracted thresholds and slopes for warm detection and heat pain were then used to determine four possible peak temperatures during the PHS task.

### PHS Stimuli

To elicit PHS, we used an adapted version of the Thermal Sensory Limen (TSL) task^23,24^, in which the peak temperature for each trial was fixed and individually determined based on thermal detection and pain thresholds. The response temperature for each trial was defined as the point at which participants reported a change in perception whilst the probe temperature was decreasing from baseline (Fig. 1). The peak temperature for each trial was set to one of four temperature intensities, generating four levels (I-1, I-2, N-1, N-2) derived from individual warm detection and heat pain PFs (Fig. 1A). I-1 corresponds to the warm detection threshold (WDT), defined as the temperature at which there is a 50% probability of detecting warmth; I-2 is the WDT plus three times the inverse of the slope of the PF. N-1 is the heat pain threshold (HPT), defined as 50% probability of detecting heat pain; and N-2 corresponds to the HPT plus two times the inverse slope. The range of possible peak temperatures was bounded between 33ºC and 50ºC to ensure safe and consistent stimulation. If the calculated values for I-1 and N-4 fell outside this range, they were adjusted to either 33ºC or 49.9ºC, respectively. For participants with shallow slopes, which lead to extreme differences between contrast conditions, an upper bound of 1.5 was used for the inverse slope. If it was not possible to define four discrete and increasing temperatures within these constraints, a default set of temperatures (34, 37, 44 and 45ºC) was used to generate contrast levels. Figure 1B shows the chosen peak temperatures for each participant. For half of the trials the probe decreased in temperature almost immediately after reaching peak temperature, whilst for the other half peak temperature was maintained at 1.5 s before the probe decreased back to baseline.

### PHS Task

Each trial began with the thermal probe increasing in temperature, at a rate of 2.5°C/sec, from baseline (32°C) to one of four peak temperatures, followed by a decrease at the same rate back to baseline. Once the probe reached baseline temperature, a tone was played to indicate that participants could respond. The probe then remained at baseline temperature for a short interval (jittered between 0.01 and 0.5 s) to decouple the alerting effects of the tone with the following temperature change. After which, the probe temperature decreased or increased at a slower rate of 1°C/sec, to allow for more precise detection of a perceptual change (Fig. 1C). Participants could only respond once the probe temperature departed from baseline. As soon as they detected a change in sensation, the participants pressed the up arrow key, which triggered the probe to return to baseline temperature. Immediately after their response a question appeared asking whether the perceived change in sensation was cold or warm. Participants responded using either the left or right arrow keys (counterbalanced across participants), and the response time between reporting a change in sensation and responding to the question was recorded. Next, participants rated their confidence in their chosen thermal quality using a VAS ranging from from “Guess” (coded as 0) to “Certain” (coded as 100). They used the left and right arrow keys to move the slider along the VAS scale, and pressed the up arrow key to confirm their confidence rating. Participants had 10 seconds to respond to each question (Fig. 1D). The task had 64 trials in total, with 16 trials per contrast condition, and lasted between 20 to 30 minutes. Each condition included 14 experimental ‘PHS’ trials, where the probe temperature continued to decrease after the tone, and two ‘catch’ trials, where the probe temperature increased after the tone. The addition of catch trials was to reduce the likelihood of participants inferring that the probe temperature was always below baseline during the critical portion of the task. Prior to the main task, participants completed eight practice trials, delivered to the same location on the non-dominant forearm. At the end of the task, participants were asked to reflect on their detection strategy and report whether they were more likely to report a change in sensation only when they were confident of their perceived thermal quality (i.e., definitely warm or definitely cold), or whether they reported changes even when they were uncertain of the temperature quality (reported in Supplementary Materials).

### Statistical analyses

Statistical analyses and visualisations were performed using Rstan^25^ in R^48^ and followed the planned pre-registration (2nd April 2024, doi.org/10.17605/OSF.IO/T5NMD). All data and code for this project are publicly available on GitHub https://github.com/Body-Pain-Perception-Lab/uncertainty-lifespan-phs and OSF https://osf.io/wxhcq.

#### Data filtering

Prior to any analyses, we implemented several control checks to ensure data quality. We aimed to exclude participants with a Michigan Neuropathy Screening Instrument (MNSI,^44,49^ score larger or equal to seven were excluded, as this is indicative of a possible undiagnosed peripheral neuropathy. No participants had an MSNI score of above 7. At the trial-level, additional data filtering was carried out to ensure that participants adhered to task instructions. Responses were excluded if either the choice or confidence response times was less than 0.25 s, as this latency is deemed too quick to reflect deliberate decision-making processes. If the detection response time was below 0.25 s then the entire trial was removed, as it is likely that the participant was reacting to the tone and not responding to a perceived change in temperature. Finally, trials with missed choice responses (> 10 s) were also removed from analysis.

Peak temperature conditions included data from trials where the peak temperature duration was both 0 and 1.5 s, as any analysis of these trials was not included in the pre-registration. In addition, For our main analysis we excluded all “catch” trials, where the probe temperature increased after the tone.

We used the “catch” trial responses to infer whether our assumptions about lapse rates for the experimental trials are correct. Across all subjects, we found no significant correlation between the total number of wrongly reported catch trials and the number of reported PHS-trials (*ρ* = -.05 (95% HDI [-0.26, 0.16]), *p* = .33), suggesting trials flagged as PHS trials are most likely correct, and not a lapse. A Bayesian mixed effects logistic regression showed that the conditional probability of an incorrect response (responding “cold”) on a “catch” trial was 0% (95 HDI [0, 1]) which was compatible with the group level estimates of the lower asymptote of psychometric function for non-catch trials which was 2.89 (95% HDI [0.5, 5.76]). The very low lapse rate for “catch” trials is indicative of a low lapse rate throughout the entire experiment.

### Multivariate modelling of PHS, choice response time and confidence

Thermal contrast was defined as the normalised temperature contrast scaled by the maximum possible temperature difference (E.q. 1). Thermal contrast was then used to inform a logistic psychometric function, defined by four parameters (Eq. 2): the threshold (*α*_*phs*_), slope (*β*_*phs*_), guess rate (*γ*_*phs*_) and lapse rate (*λ*_*phs*_). This psychometric curve estimates the latent probability of PHS (*p*_*phs*_) as a function of thermal contrast (TC) for every participant.

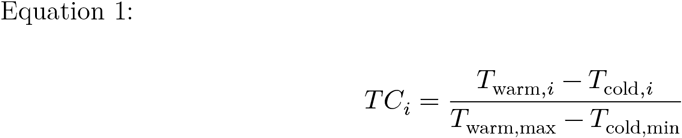

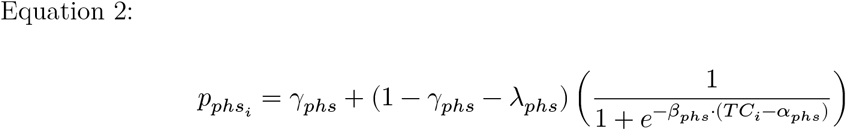

To further categorise the perceptual decision making of the PHS experience, we included response times and confidence ratings into the same model. This inclusion of additional behavioural responses quantified important aspects of the PHS experience that would be missed by only looking at the binary responses. Using all three behavioural measures we developed two multivariate models, based around two possible PHS profiles (true perceiver and unsure perceiver), with the main difference of how uncertainty about the PHS experience is affecting the response times and confidence ratings (Fig. 3). We assume that conditions that consistently lead to longer response times and lower reported confidence are suggestive of a greater uncertainty in choosing between “cold” and “warm”. This pattern can be used to determine whether specific stimulus conditions contribute to more ambiguous sensory experiences typically associated with PHS.

#### Model 1: True perceiver

The true perceiver assumes that perceptual uncertainty in PHS arises from the difficulty of making a categorical choice between “cold” and “warm”, rather than from the experience of PHS itself. We refer to this profile as the true perceiver, because certainty is highest at the points at which the sensation is either clearly cold, or clearly warm (i.e., PHS as a seemingly veridical warm sensation). Therefore, this model predicts the fastest response times and highest confidence at thermal contrast values where participants either feel veridical cold or report PHS with near certainty. In contrast, this model predicts the slowest response times and lowest confidence at thermal contrast values associated with 50% probability of experiencing a PHS, where the stimulus is most ambiguous. This pattern aligns with the concept of a “true” PHS responder as an individual who experiences a clear and definite warm sensation in response to a cold stimulus.

For the true perceiver model we quantify uncertainty as Shannon entropy of the probability of responding PHS (Eq. S1.1), which is linked to decision difficulty. We map the Shannon entropy to both response time and confidence ratings as affine functions.

#### Model 2: Unsure perceiver

The unsure perceiver assumes that perceptual uncertainty in PHS arises not from the difficulty of making a categorical choice between “cold” and “warm”, but from the aberrant and unusual experience of PHS, therefore uncertainty increases with the probability of experiencing PHS itself. This profile is called the unsure perceiver as participants are more uncertain about their perception at stimulus intensities where participants more frequently report PHS, reflected in longer response times and lower confidence ratings. Thus for the unsure perceiver we map the probability of responding PHS directly to the response times and confidence ratings through affine functions.

Both true and unsure perceiver models response times and confidence ratings were estimated with the addition of thermal contrast as a regressor, in order to disentangle the contribution of the reporting of PHS from the thermal contrast itself (see Supplementary Materials for a full set of equations and description of the full models).

#### Model Estimation and Selection

All models were fitted using Stan’s Hamiltonian Markov chain Monte Carlo No U-turn sampler^25^. To ensure that all models fit our defined diagnostic criteria, we increased the target average acceptance probability from .95 to .99 and the maximum tree depth from 12 to 15 to ensure no divergent transitions, as outlined in the contingency plans of the pre-registered analyses. We also increased the number of warm-up and post warm-up samples from 1,000 to 2,000 for each of the four chains to ensure R-hat values below 1.01 and effective sample size above 400 diagnostic values. Further, we inspected trace and pair plots to investigate sampling of the parameter space. We specified hyperpriors based on simulated data and prior knowledge^23^.

Model selection was conducted using approximate leave-one-out cross-validation, as implemented in the loo package^50^. We initially applied Pareto smoothed importance sampling leave-trial-out cross-validation. However, both models had Pareto-k’s (diagnostic measure) exceeding 0.7, (0.73%) and (0.71%) for the true perceiver and unsure perceiver model, respectively. In an attempt to address this, we switched to moment matching importance sampling leave-trial-out cross-validation, but the same trials remained problematic. Model performance was evaluated using ELPD, with higher values indicating better ability to explain the data.

We tested the two multivariate models through parameter recovery to ensure both could accurately recover their generating parameters, and also performed model recovery to ensure discriminability between the two generative models. We further used these simulations to investigate the feasibility of including an age regressor on both threshold and slope of the psychometric function, which revealed that we were only able to detect a difference in the threshold given the assumed effect sizes. Full details are documented in the pre-registration and Fig. S6.

## Acknowledgements

We would like to thank Johanne Sejrskild Rejsenhus for her help with data collection and Dr. Camila Sardeto Deolindo for her contribution to the thermal thresholding procedure.

## Author contributions

Conceptualisation: J.F.E., A.G.M. & F.F. Methodology: A.G.M., C.E.K. & F.F. Investigation: J.F.E., A.G.M., C.E.K., A.S.C. & F.F. Data analysis: J.F.E., A.G.M., A.S.C. & C.S. Visualisation: J.F.E., A.G.M. & F.F. Writing - original draft: J.F.E., A.G.M. & F.F. Writing - review & editing: J.F.E., A.G.M., C.E.K., A.S.C.& F.F. Supervision: F.F.

