## Supplementary Materials for "Modelling perceptual uncertainty in a thermosensory illusion across the lifespan"

### Deviations from pre-registration

The pre-registration inaccurately reported the total number of participants as 83, this information was corrected in the manuscript: N = 84 in total and N = 75 after exclusions.

### Additional results

#### PHS prevalence

Figure S1 shows the total number of participants that experienced a PHS, and the max number of PHS per participant for each peak temperature condition.

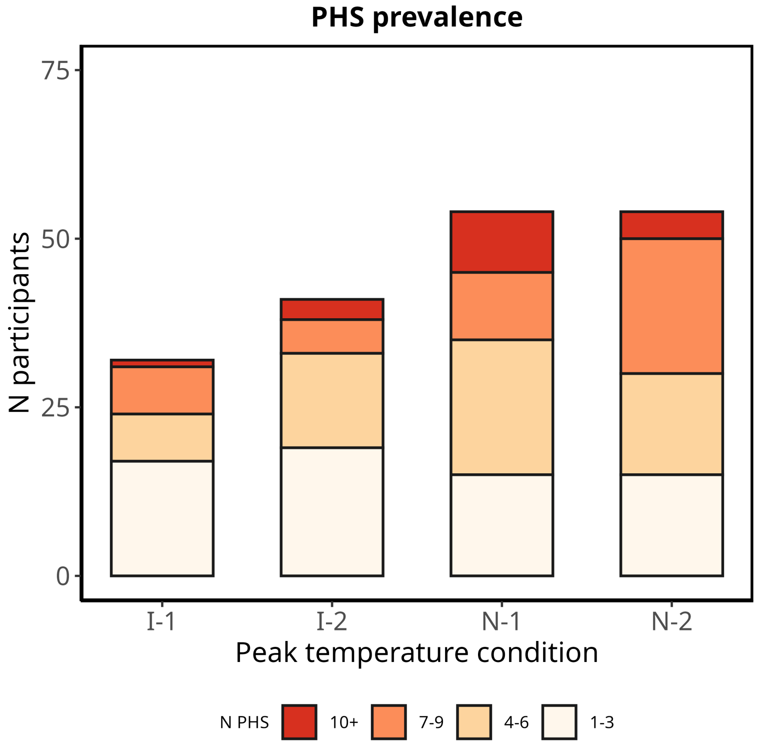


**Figure S1:** Histogram of the number of PHS responders (i.e., participants reporting at least one PHS) for each peak temperature condition. Shaded colours show the total number of PHS reported (binned) for those participants.

### Peak temperatures

To determine whether the age-related increase in thermal contrast was driven by the warming or cooling phase of the task, we tested whether peak temperatures varied with age. Linear regression showed peak temperatures increased with increasing age across all conditions ($\beta = 0.03$, 95% HDI [0, 0.06], $p = .97$. However, there was no effect of age on peak temperatures within each temperature condition.

### Task-related self reports on response strategy

At the very end of the task, we asked participants to reflect on their response strategy. First, they indicated whether they reported a change in perceived temperature when they were unsure, or whether they waited until they were sure of their perceived temperature quality (forced-choice question). Following this, they were asked to rate, on two, sequential VAS scale from 0 to 100, how often they reported a change in temperature when they were (1) unsure and (2) sure of the sensation they were perceiving.

In our final sample of 75 participants, 24 (32%) reported pressing a button to indicate a change in sensation when they were unsure of the temperature quality, whilst 51 (68%) waited until they were sure. Forced-choice responses were not influenced by age ($\beta = 0.01$, 95% HDI [-0.01, 0.04], $p = .87$). However, task-related self reports changed slightly with age. An ordered-beta mixed-effect regressions indicated that "sure" VAS ratings increased across the lifespan ($\beta = 0.01$, 95% HDI [0, 0.03], $p = .98$), suggesting that older adults were more likely to wait until they were confident about the thermal quality before reporting a temperature change. No significant effect of age was observed on "unsure" VAS ratings ($\beta = 0.01$, 95% HDI [0, 0.02], $p = .93$). The age-related increase in "sure" ratings may help explain the increased confidence among older adults. However, this could also reflect a response bias, with older participants tending to use the upper end of the rating scale more frequently.

### Responses and posterior predictive checks from all participants

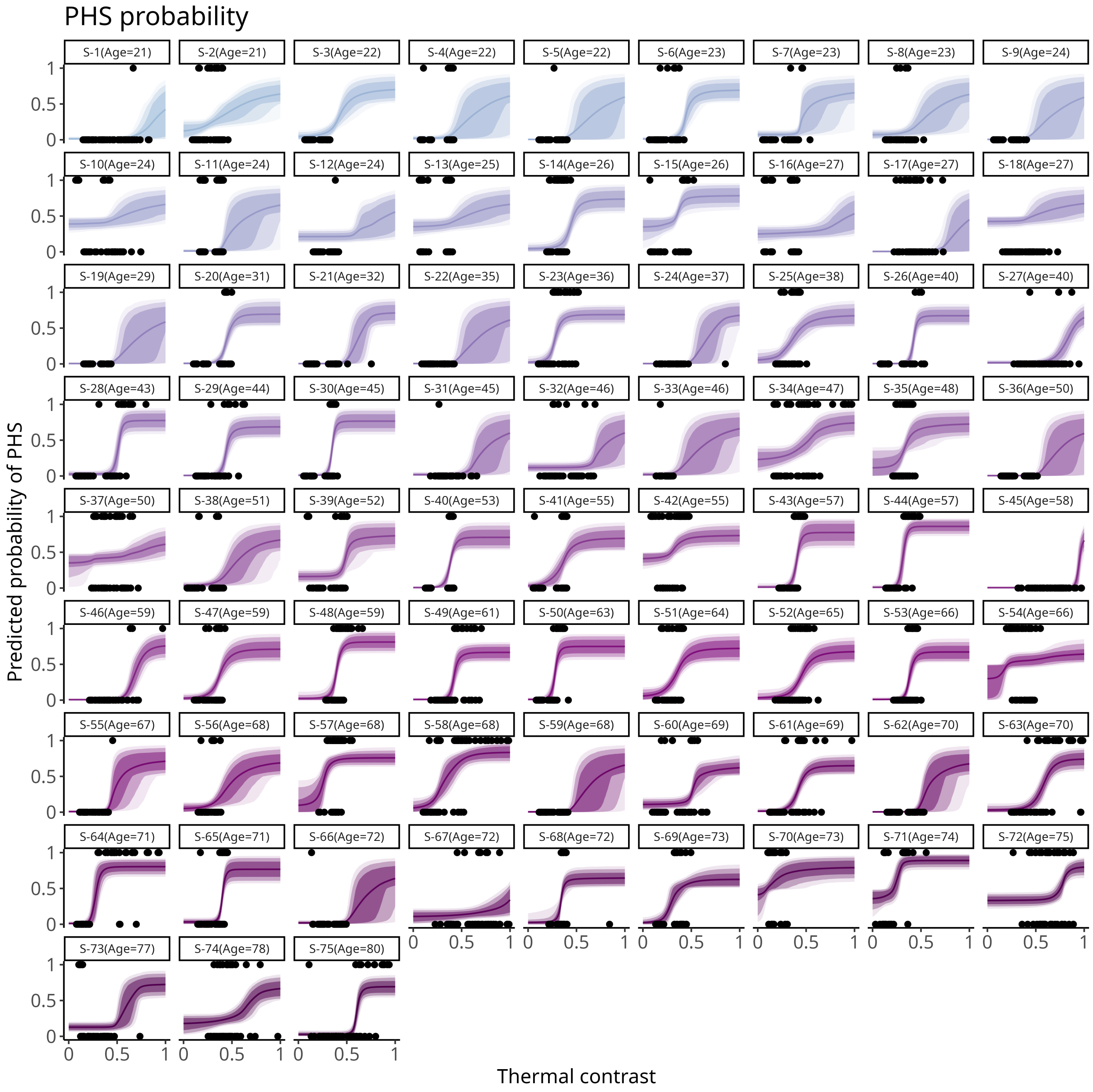


**Figure S2:** Binary responses and posterior estimates of predicted PHS probability as a function of thermal contrast for each participant, ordered by age. Solid line shows the mean estimate, banded error bars show 95%, 90% and 80% credible intervals.

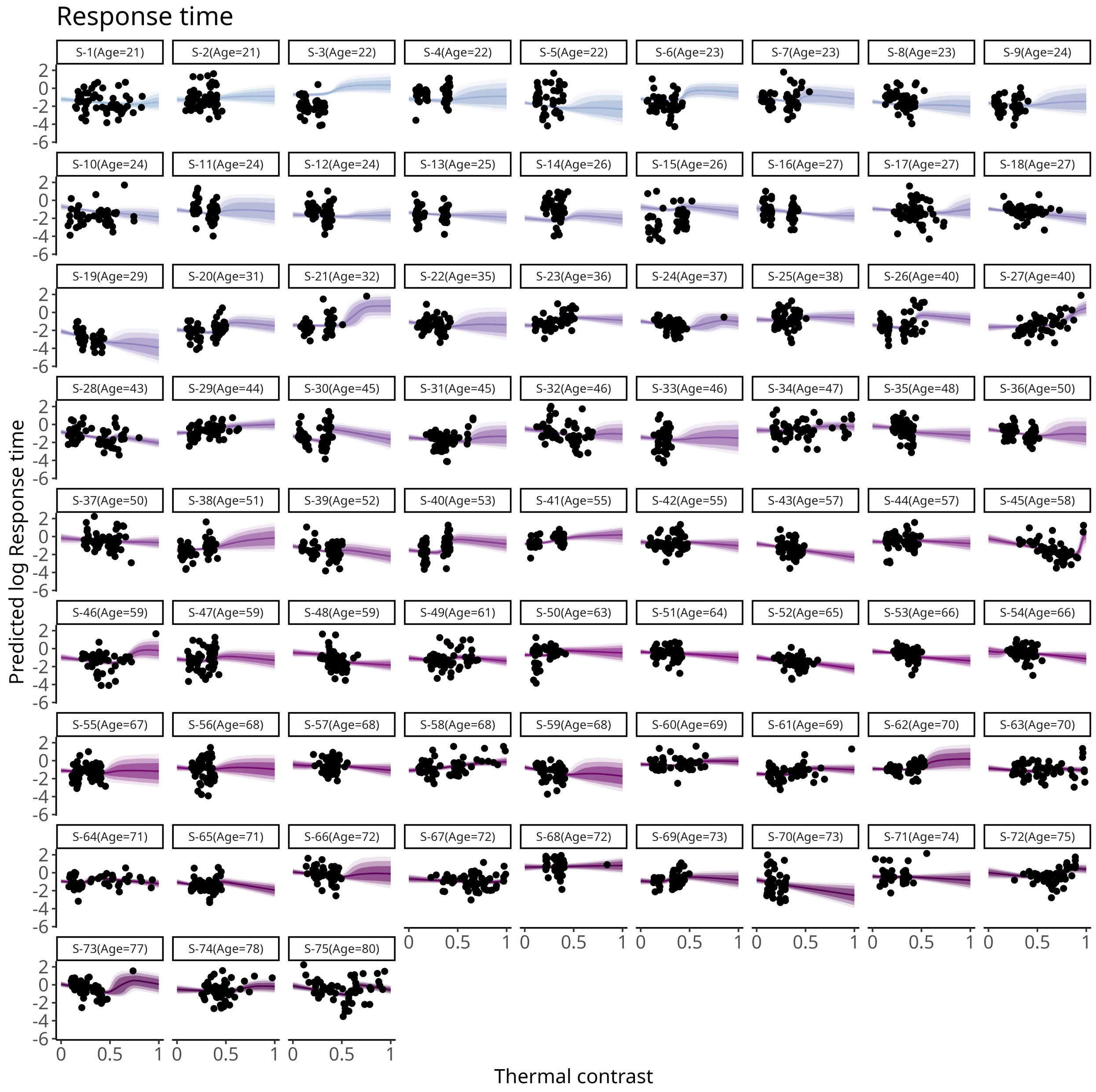


**Figure S3:** Log response times and posterior estimates of mean log response time as a function of PHS probability and thermal contrast for each participant, ordered by age. Solid line shows the mean estimate, banded error bars show 95%, 90% and 80% credible intervals.

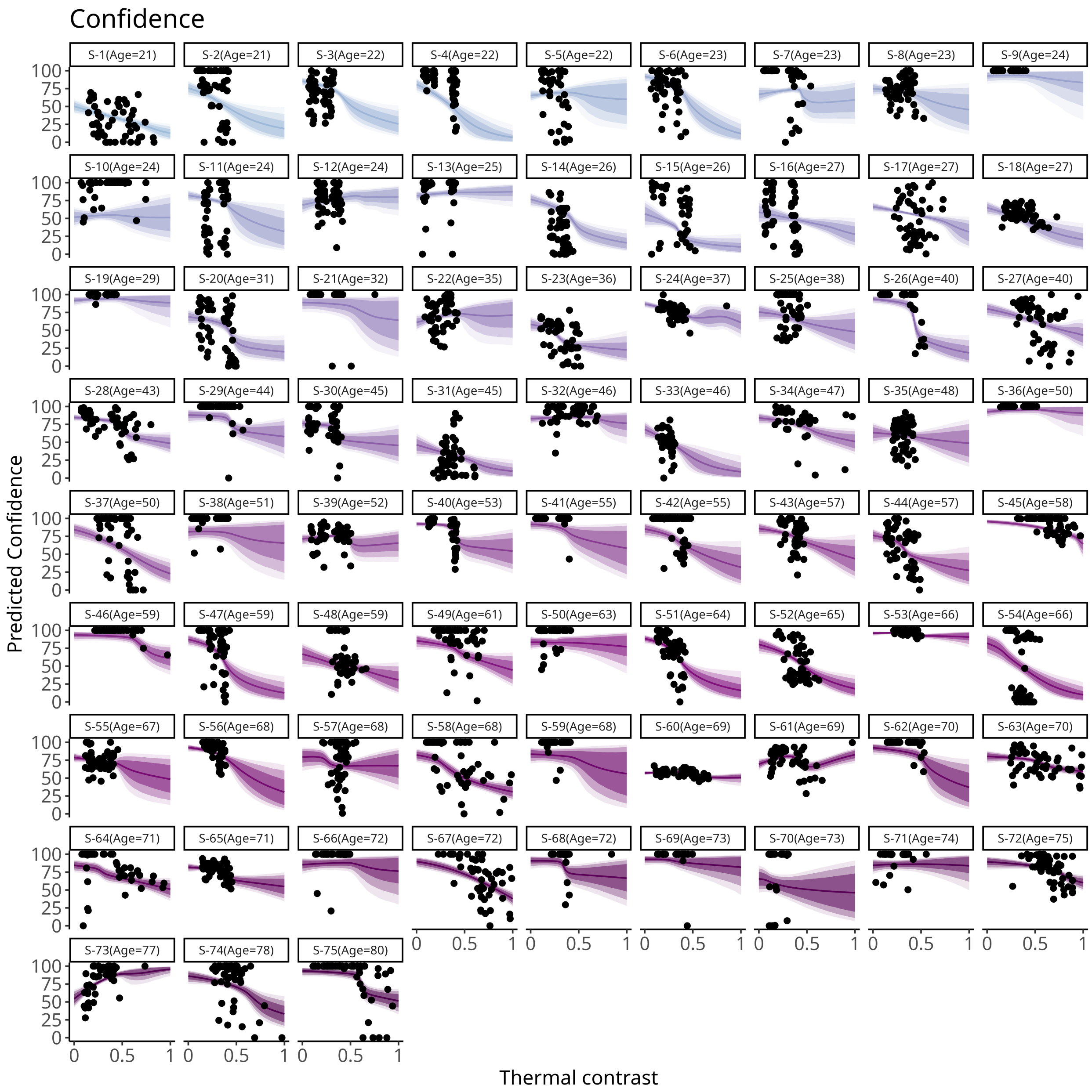


**Figure S4:** Confidence ratings and posterior estimates of mean confidence as a function of PHS probability and thermal contrast for each participant, ordered by age. Solid line shows the mean estimate, banded error bars show 95%, 90% and 80% credible intervals.

### Confidence outlier

Figure 6 from the manuscript, including outlier with inflated estimated effect of thermal contrast on confidence

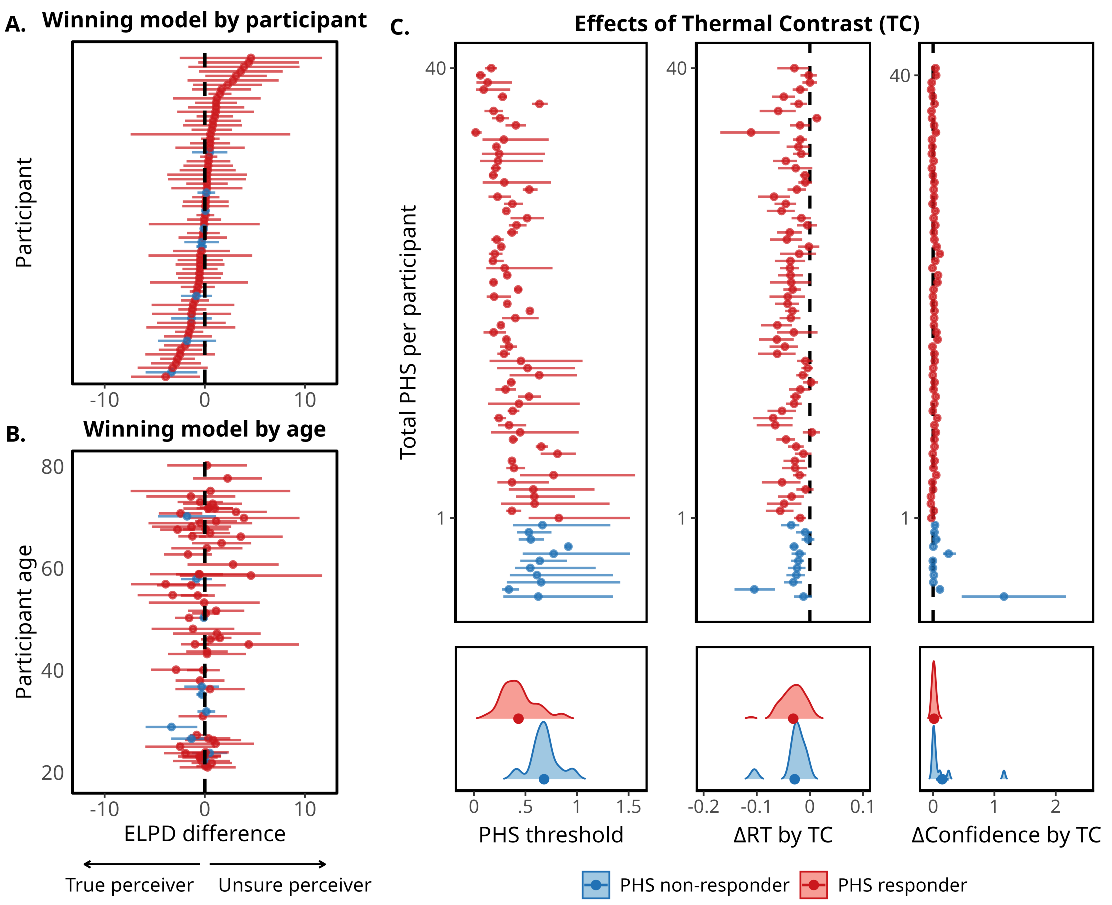


**Figure S5:** A reproduction of Figure 6 from the main manuscript, with the outlier included in the Confidence estimates in panel C.

### Model Description and Priors

#### Model 1: True perceiver

In the true perceiver model, we assume that uncertainty scales with the Shannon entropy of the binary decision process, defined as:

**Equation S1.1**

$$H(phs) = -\left(p_{phs} \cdot \log(p_{phs}) + (1 - p_{phs}) \cdot \log(1 - p_{phs})\right)$$

This operationalization of uncertainty is then linearly linked to both the mean response time and confidence rating together with two additional terms; an intercept ($\alpha$), and a term for the influence of thermal contrast beyond the uncertainty introduced by the probability of PHS ($\beta_2$).

**Equation S1.2 (Response times)**

$$rt_t = \alpha_{rt_s} + b1_{rt_s} \cdot H(p_{phs_t}) + b_{2rt_s} \cdot TC_t + b_{rt_{age}} \cdot Age$$

**Equation S1.3 (Confidence)**

$$conf_t = \alpha_{conf_s} + b1_{conf_s} \cdot H(p_{phs_t}) + b_{2conf} \cdot TC_t + b_{conf_{age}} \cdot Age$$

#### Model 2: Unsure perceiver

In the unsure perceiver model, we operationalised uncertainty as the probability of PHS, defined as:.

**Equation S1.4 (Response time)**

$$rt_t = \alpha_{rt_s} + b1_{rt_s} \cdot p_{phs_t} + b_{2rt_s} \cdot TC_t + b_{rt_{age}} \cdot Age$$

**Equation S1.5 (Confidence)**

$$conf_t = \alpha_{conf_s} + b1_{conf_s} \cdot p_{phs_t} + b_{2conf} \cdot TC_t + b_{conf_{age}} \cdot Age$$

Note that both models included a between-subject parameter for the effect of age on the threshold of the psychometric function.

**Equation S1.6**

$$\alpha_{PHS} = \alpha_\beta + b_{PHS_{age}} \cdot Age$$

### Likelihood Functions

The probability of experiencing PHS were mapped to the participant's binary responses using the Bernoulli likelihood distribution (Eq. S1.7). Response times were fitted using a shifted log-normal likelihood function with a participant-specific mean ($\mu_{rt}$), standard deviation ($\sigma_{rt}$), and a non-decision time ($\tau$, Eq. S1.8). To model confidence ratings, we used the ordered beta likelihood, which combines both an ordinal and beta regressions to fully capture data with both upper and lower bounds (e.g., VAS ratings, Eq. S1.9). The parameters of the model contain the mean confidence ($\mu_{conf}$), a precision estimate ($\phi_{conf}$) and two defined cut-points to represent distribution of 0s ($c_1$) and 1s ($c_1$) in the data.

**Equation S1.7**

$$PHS \sim \text{Bernoulli}(phs)$$

**Equation S1.8**

$$(RT + \tau) \sim \text{LogNormal}(rt, \sigma_{rt})$$

**Equation S1.9**

$$Confidence \sim \text{OrderedBeta}(\mu_{conf}, \phi_{conf}, c1, c2)$$

The residual variance in the shifted-lognormal and precision parameter in the ordered beta likelihoods were exponentiated to ensure their positive constraint. Further, the non-decision time was constrained between 200 ms and the minimum response time for that subject. The two cut-points for the ordered beta likelihood were constrained so that the lower cut-point was smaller than the upper, achieved using an induced Dirichlet prior. Lastly, for the parameters of the psychometric function, we constrained the lapse and guess rates (lower and upper asymptote) between 0 and 0.5. This parameterisation together with a strictly positive slope parameter ensured that the model was identifiable (i.e., can not have two solutions). Lastly, we exponentiated our threshold priors, constrained it to be positive in order to simulate data that adhered to expectation (i.e. a long tailed distribution for the psychometric threshold which captures potential non-responders). These decisions were made prior to pre-registration.

### Priors

Here we present prior values used to fit models 1 and 2.

#### Group-level priors

Psychometric function:

$$\mu_{\beta_{PHS}} \sim N(-2, 1.5), \quad \sigma_{\beta_{PHS}} \sim N(0, 1), \quad \mu_{\beta_{\alpha_{PHS}}} \sim N(4, 0.5), \quad \sigma_{\beta_{\alpha_{PHS}}} \sim N(1, 0.5)$$

$$\mu_{\gamma_{PHS}} \sim N(-5, 1), \quad \sigma_{\gamma_{PHS}} \sim N(0, 1), \quad \mu_{\beta_{\lambda_{PHS}}} \sim N(0, 1), \quad \sigma_{\beta_{\lambda_{PHS}}} \sim N(0, 1)$$

Response time effects:

$$\mu_{\alpha_{rt}} \sim N(0.3, 0.5), \quad \sigma_{\alpha_{rt}} \sim N(1, 1), \quad \mu_{\beta_{1rt}} \sim N(2.5, 1), \quad \sigma_{\beta_{1rt}} \sim N(1, 2)$$

$$\mu_{\beta_{2rt}} \sim N(0, 1), \quad \sigma_{\beta_{2rt}} \sim N(1, 2), \quad \mu_{\tau} \sim N(-2, 1), \quad \sigma_{\tau} \sim N(0, 1),$$

$$\mu_{\sigma_{rt}} \sim N(-1, 0.5), \quad \sigma_{\sigma rt} \sim N(0.5, 1)$$

Confidence effects:

$$\mu_{\alpha_{conf}} \sim N(0.3, 0.5), \quad \sigma_{\alpha_{conf}} \sim N(1, 1), \quad \mu_{\beta_{1conf}} \sim N(2.5, 1), \quad \sigma_{\beta_{1conf}} \sim N(1, 2)$$

$$\mu_{\beta_{2conf}} \sim N(0, 1), \quad \sigma_{\beta_{2conf}} \sim N(1, 2), \quad \mu_{c1} \sim N(-2, 1), \quad \sigma_{c1} \sim N(0, 1)$$

$$\mu_{c2} \sim N(2, 0.25), \quad \sigma_{c2} \sim N(0, 1), \quad \mu_{\phi_{conf}} \sim N(3, 0.5), \quad \sigma_{\phi_{conf}} \sim N(0, 1)$$

Age-related effects:

$$(\beta_{rt_{age}}, \quad \beta_{rt_{age}}, \quad \beta_{rt_{age}}) \sim N(0, 0.3)$$

**Subject-level priors**

**Psychometric function parameters**

$$\beta_{PHS_s} \sim \exp\left(\mathcal{N}(\mu_{\beta_{PHS}}, \sigma_{\beta_{PHS}})\right), \quad \beta_{\alpha_{PHS_s}} \sim \exp\left(\mathcal{N}(\mu_{\beta_{\alpha_{PHS}}}, \sigma_{\beta_{\alpha_{PHS}}})\right)$$

$$\gamma_{PHS_s} \sim S^{-1}\left(\mathcal{N}(\mu_{\gamma_{PHS}}, \sigma_{\gamma_{PHS}})\right), \quad \beta_{\lambda_{PHS_s}} \sim S^{-1}\left(\mathcal{N}(\mu_{\beta_{\lambda_{PHS}}}, \sigma_{\beta_{\lambda_{PHS}}})\right)$$

**Response time parameters**

$$\alpha_{rt_s} \sim \exp\left(\mathcal{N}(\mu_{\alpha_{rt}}, \sigma_{\alpha_{rt}})\right), \quad \beta_{1rt_s} \sim \mathcal{N}(\mu_{\beta_{1rt}}, \sigma_{\beta_{1rt}})$$

$$\beta_{2rt_s} \sim \mathcal{N}(\mu_{\beta_{2rt}}, \sigma_{\beta_{2rt}}), \quad \tau_s \sim S^{-1}\left(\mathcal{N}(\mu_\tau, \sigma_\tau)\right)$$

$$\sigma_{rt_s} \sim \exp\left(\mathcal{N}(\mu_{\sigma_{rt}}, \sigma_{\sigma_{rt}})\right)$$

**Confidence parameters**

$$\alpha_{conf_s} \sim \mathcal{N}(\mu_{\alpha_{conf}}, \sigma_{\alpha_{conf}}), \quad \beta_{1conf_s} \sim \mathcal{N}(\mu_{\beta_{1conf}}, \sigma_{\beta_{1conf}})$$

$$\beta_{2conf_s} \sim \mathcal{N}(\mu_{\beta_{2conf}}, \sigma_{\beta_{2conf}}), \quad \phi_{conf_s} \sim \exp\left(\mathcal{N}(\mu_{\phi_{conf}}, \sigma_{\phi_{conf}})\right)$$

$$c1_s \sim \mathcal{N}(\mu_{c1}, \sigma_{c1}), \quad c2_s \sim \mathcal{N}(\mu_{c2}, \sigma_{c2})$$
